# Aperiodic brain activity and response to anesthesia vary in disorders of consciousness

**DOI:** 10.1101/2022.04.22.489199

**Authors:** Charlotte Maschke, Catherine Duclos, Adrian M. Owen, Karim Jerbi, Stefanie Blain-Moraes

**Affiliations:** Montreal General Hospital, McGill University Health Centre, Montreal, Canada; Integrated Program in Neuroscience, McGill University, Montreal, Canada; Hôpital du Sacré-Cœur de Montréal, Centre intégré universitaire de Santé et de Services Sociaux du Nord-de-l’île-de-Montréal, Montréal, Québec Canada; Department of Anesthesiology and Pain Medicine, Université de Montréal, Montréal, Québec Canada; Department of Physiology and Pharmacology, Western University, London, Ontario, Canada; Western Institute for Neuroscience, Western University, London, Ontario, Canada; Department of Psychology, Western University, London, Ontario, Canada; Cognitive & Computational Neuroscience Lab, Psychology Department, University of Montreal, Québec, Canada, MILA (Québec Artificial Intelligence Institute), Montréal, Québec, Canada; Centre UNIQUE (Union Neurosciences & Intelligence Artificielle), Montréal, Québec, Canada; School of Physical and Occupational Therapy, McGill University, Montreal, Canada

## Abstract

The analysis of human EEG has traditionally focused on oscillatory power, which is characterized by peaks above an aperiodic component in the power spectral density. This study investigates the aperiodic EEG component of individuals in a disorder of consciousness (DOC); how it changes in response to exposure to anesthesia; and how it relates to the brain’s information richness and criticality. High-density EEG was recorded from 43 individuals in a DOC, with 16 of these individuals undergoing a protocol of propofol anesthesia. The aperiodic component was defined by the spectral slope of the power spectral density. Our results demonstrate that the EEG aperiodic component is more informative about the participants’ level of consciousness than the oscillatory component. Importantly, the pharmacologically induced change in the spectral slope from 30-45 Hz positively correlated with individual’s pre-anesthetic level of consciousness. The pharmacologically induced loss of information-richness and criticality was associated with individual’s pre-anesthetic aperiodic component. During exposure to anesthesia, the aperiodic component was correlated with 3-month recovery status for individuals with DOC. The aperiodic EEG component has been historically neglected; this research highlights the necessity of considering this measure for the assessment of individuals in DOC and future research that seeks to understand the neurophysiological underpinnings of consciousness.

## Introduction

Are there signatures in human brain activity that can be used to delineate an individual’s state of consciousness? In the quest for electrophysiological markers of consciousness, the analysis of the human electroencephalogram (EEG) has traditionally been focused on oscillatory patterns within specific frequency bands, which are typically present as peaks in the power-spectral density (PSD) of the human EEG. However, oscillatory peaks always co-occur with broadband non-oscillatory (i.e. aperiodic) activity, which can be described by the exponential (i.e. 1/f-like) decay of power over frequency (Donoghue *et al*., 2020; Donoghue, Schaworonkow and Voytek, 2021). Recent advances in electrophysiology (Donoghue *et al*., 2020; Donoghue, Schaworonkow and Voytek, 2021) suggest that analyzing EEG data solely from the perspective of oscillatory patterns may lead to erroneous or incomplete representations of the underlying neurophysiological processes.

Electroencephalography is a particularly promising tool for assessing the level of consciousness of individuals in a disorder of consciousness (DOC), as it is highly accessible in the clinical setting, has few patient contraindications and can be recorded at the bedside (Swisher and Sinha, 2016). Individuals in a DOC following brain injury exhibit a wide range of reduced levels of awareness and arousal. As consciousness and responsiveness can be completely dissociated (Owen *et al*., 2006; Sanders *et al*., 2012; Mashour and Avidan, 2013), the identification of behavior-independent measures of consciousness is crucial for uncovering the mechanisms of human consciousness and improving clinical practice.

Although the PSD of healthy adult EEG is characterized by the presence of spectral peaks — predominantly in the theta and alpha bandwidth — the PSD of individuals in DOC often exhibits a total absence of such peaks (see Fig 1A). Interpreting the remaining power of the aperiodic component erroneously as being oscillatory leads to several significant methodological problems (Donoghue, Schaworonkow and Voytek, 2021). When not considered separately, putative changes in EEG oscillations across tasks and conditions might be underpinned entirely by alterations in the aperiodic component of the EEG. Thus, investigating the aperiodic component in the EEG of individuals in DOC might also lead to more complete representations of the neurophysiological underpinnings of consciousness.

**Figure 1.**
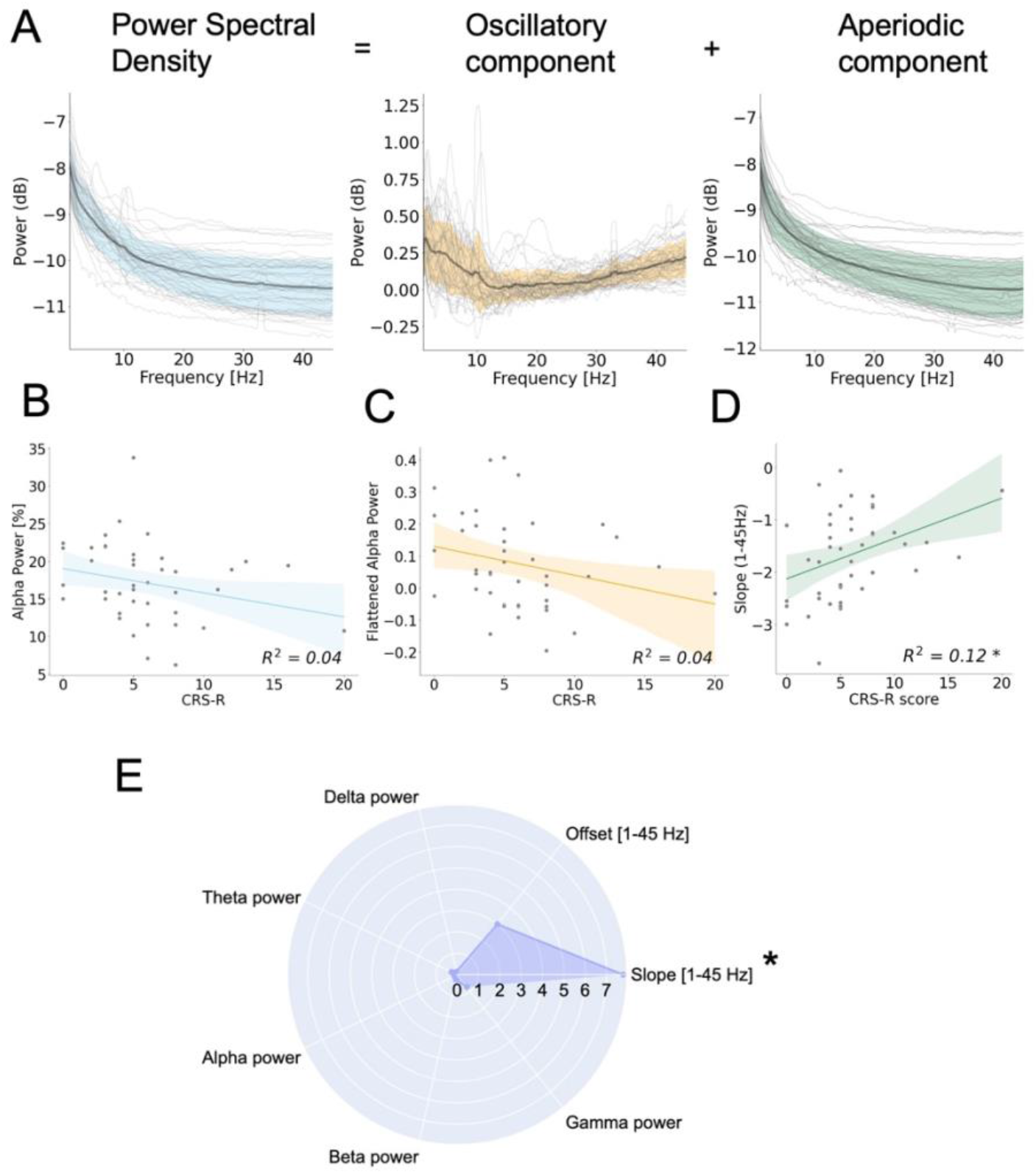
Diagnostic value of EEG spectral properties in DOC. **(A)** The power spectral density of individuals in DOC (blue) separates into an oscillatory component (orange) and an aperiodic component (green). Light grey lines represent individual subjects; darker lines represent the group average and standard deviation. **(B)** The traditional power analysis of oscillatory EEG in the alpha bandwidth does not predict individual’s level of consciousness, as measured by the CRS-R score.^1^ **(C)** After removal of the aperiodic component, remaining oscillatory power of the EEG in the alpha bandwidth does not predict individuals’ level of consciousness, as measured by the CRS-R score.^1^ **(D)** The aperiodic component of individuals EEG predicts individual level of consciousness. ^1^ **(E)** Model coefficients after combining oscillatory and aperiodic features in multivariate linear regression model. Only the aperiodic slope is a significant predictor for individuals’ level of consciousness. ^1^ for the purpose of visualization, R^2^ were obtained through univariate linear regressions. * indicates *p*<0.05.

Recent studies have shown that the properties of the aperiodic EEG contain information about consciousness, which are neglected in traditional oscillation-based analyses (Colombo *et al*., 2019; Lendner *et al*., 2020). States of unconsciousness, such as non-rapid eye movement sleep (Lendner *et al*., 2020) and anesthetic-induced unconsciousness (Colombo *et al*., 2019; Lendner *et al*., 2020) exhibit a steeper spectral slope (i.e. a faster power decay over frequencies), compared to wakefulness. Changes in the aperiodic EEG were further observed after exposure to psychoactive drugs (Muthukumaraswamy and Liley, 2018; Timmermann *et al*., 2019). As such, the aperiodic component has been widely proposed as an electrophysiological marker for the assessment of individuals in a DOC (Colombo *et al*., 2019; Lendner *et al*., 2020).

During conscious wakefulness, the brain has been widely suggested to operate close to criticality — a point of optimal computational capacity and information-richness where the underlying network is poised between order and disorder ((Beggs and Plenz, 2003; Carhart-Harris *et al*., 2014; Carhart-Harris, 2018; Zimmern, 2020; O’Byrne and Jerbi, 2022; Toker *et al*., 2022). This balance is putatively maintained by a proper tuning between excitation and inhibition (E/I) (Shew et al., 2011; Zhou and Yu, 2018). The aperiodic EEG and has been linked to the local E/I balance, the brain’s information-richness and criticality (Gao, Peterson and Voytek, 2017; Muthukumaraswamy and Liley, 2018; Medel *et al*., 2020; Toker *et al*., 2022). Being additionally correlated to established consciousness metrics, such as the Perturbation Complexity Index (Casali *et al*., 2013; Colombo *et al*., 2019), the aperiodic component has shown much promise for the investigation of mechanisms underlying consciousness, especially in individuals who are behaviorally unresponsive.

Using EEG recorded under various conditions of pharmacologically and pathologically induced unconsciousness, this study aimed to characterize the aperiodic component associated with consciousness for individuals in DOC, and in particular, how this component changes in response to exposure to anesthesia and how it relates to altered network complexity and criticality. General anesthesia is known to reliably reduce levels of consciousness *and* responsiveness by globally perturbing brain networks underlying consciousness (Purdon *et al*., 2013). Investigating the anesthetic-induced change of the aperiodic component in individuals in DOC provides a unique perspective on mechanisms underlying human consciousness. We first hypothesized that the aperiodic EEG component would have diagnostic value for individuals in DOC above and beyond the traditional analysis of EEG oscillatory power. We further hypothesized that the pharmacologically induced change in the aperiodic component would vary with individuals’ level of, and capacity for, consciousness, and that this change would be accompanied by the brain’s loss of network criticality.

## Results

This study combined two existing datasets of DOC participants (n= 43), a subset of whom (n= 16) were exposed to a targeted infusion of propofol anesthesia (see Methods). To characterize the aperiodic EEG component associated with level of consciousness, we first analyzed an existing dataset of 128-channel EEG recorded from 43 individuals in DOC at resting state. As a surrogate for consciousness, the level of responsiveness of each participant was assessed by a trained experimenter using the Coma Recovery Scale-Revised (CRS-R) (Kalmar and Giacino, 2005). For participants in an acute DOC (n = 18), recovery of consciousness was assessed three months post-EEG. At this time, six participants had recovered full consciousness (i.e., were able to respond verbally and consistently follow commands) (see Methods). The PSD was calculated using the Multitaper method. The aperiodic component of the EEG was defined by the offset and slope of the PSD (i.e., a steeper slope indicating faster decay of power over frequencies). Both parameters of the aperiodic component were estimated from 1-45 Hz using the ‘Fitting oscillations and one over f’ (FOOOF) algorithm (Donoghue *et al*., 2020) (see Methods). Oscillatory power in the delta (1-4 Hz), theta (4-8 Hz), alpha (8-13 Hz), beta (13-30 Hz) and gamma (30-45 Hz) bandwidth was calculated before and after the removal of the aperiodic component. The detection of oscillatory peak frequency was performed using the FOOOF algorithm.

### Aperiodic EEG component contains more diagnostic value than oscillatory power for individuals in a DOC

We conducted a multiple regression analysis to investigate the diagnostic information of the aperiodic component over and above the traditional power analysis. We combined oscillatory power over five canonical frequency bands (i.e., delta, theta, alpha, beta, gamma without removing the aperiodic component) and aperiodic features (i.e., spectral slope and offset) in one multivariate linear regression model.

The spectral slope was the only significant predictor of participants’ CRS-R score (see Table 1) and explained 12% of the variance (R^2^=0.12, F(1,40)=6.67, p<0.05) (see Fig. 1D). Individuals with higher levels of consciousness exhibited a flatter slope (see Fig. 1A). In contrast to previous research (Lechinger *et al*., 2013; Chennu *et al*., 2014), we did not find a significant relation between spectral power in any frequency band and participants’ CRS-R score (see Fig.1B, Fig.S1 for all frequency bands). The oscillatory-only component (i.e., after removal of the aperiodic component) was not predictive for an individual’s CRS-R score (see Fig. 1C, Fig. S1 for all frequency bands).

**Table 1.**
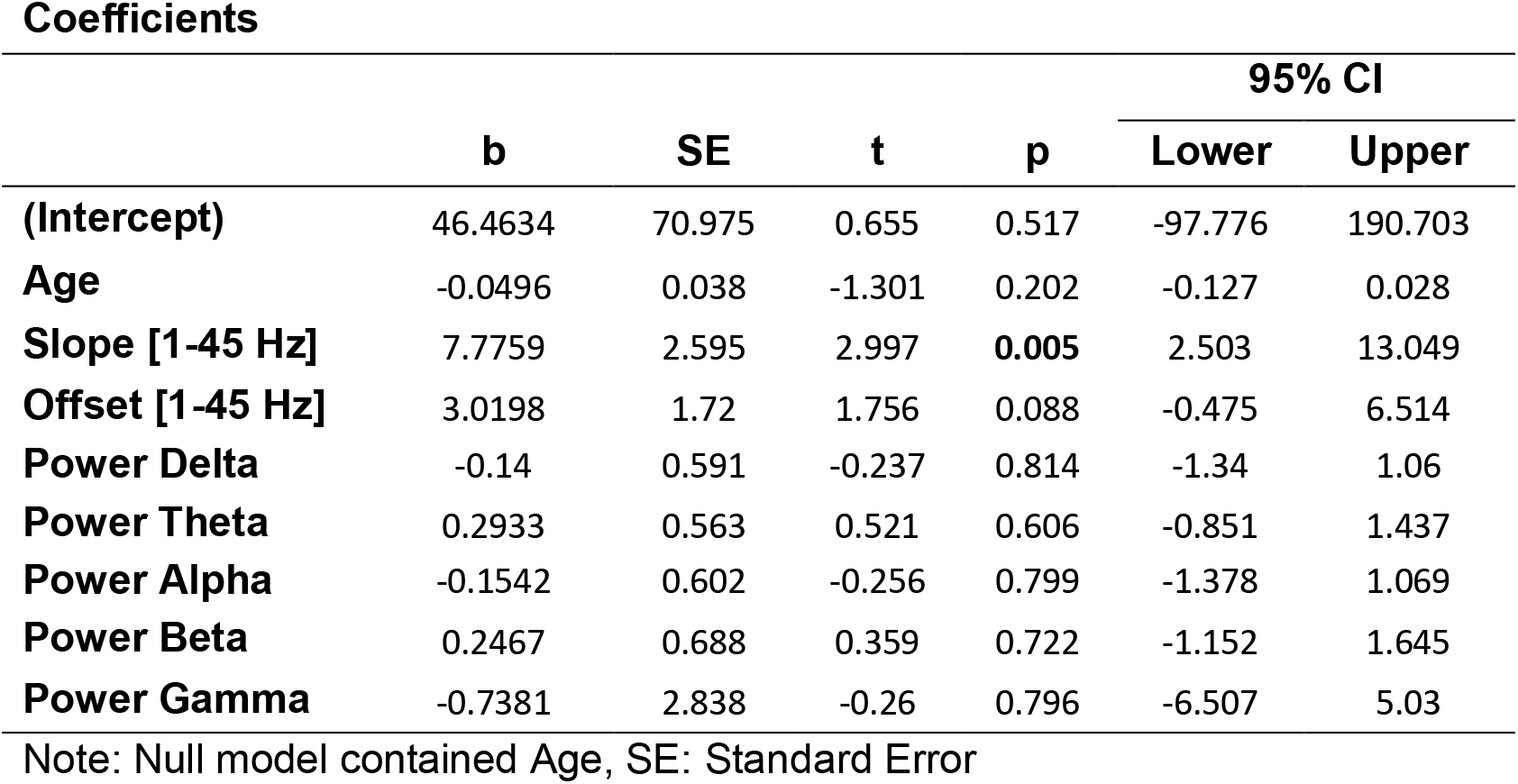
Linear model with combined band-limited power and aperiodic features

An oscillatory peak could be identified in only 13 out of 43 individuals in DOC (see Fig. S2). Peaks were predominantly identified in the theta to lower-alpha frequency range (5.1 Hz ± 3.01). Neither the peak frequency, nor the power of the identified peak oscillation (for n = 13) correlated with individuals’ CRS-R score (see Fig. S2, see Discussion). Most importantly, no oscillatory peak was detected in the remaining 30 participants, leaving solely the aperiodic EEG component for analysis.

To control for known changes of the aperiodic component over lifespan (Voytek *et al*., 2015), all models were controlled for participants’ age. There was no significant correlation between the offset of the aperiodic component and participants’ CRS-R score. The spectral slope was significantly steeper in acute DOC, compared to individuals with chronic DOC (t(41) = 2.63, p=0.01) (see Fig. S1). There was no significant difference in the spectral slope or the identified peak between participants who did or did not recover consciousness 3 months following EEG assessment. In this analysis, the aperiodic component was estimated using the ‘knee mode’ of the FOOOF algorithm (min_peak_height=0.1, max_n_peaks=10) (Donoghue *et al*., 2020), which led to a smaller model error compared to the ‘fixed mode’; all results were replicated using the ‘fixed mode’ (see Fig. S1).

### Participants with higher levels of consciousness exhibit larger changes of the aperiodic EEG component in response to anesthesia

Measuring the brain’s reaction to perturbations has shown strong potential for the assessment of individuals in altered states of consciousness (Casali *et al*., 2013; Duclos *et al*., 2022) (see Discussion). General anesthesia is known to globally perturb brain networks, resulting in loss of consciousness *and* responsiveness. We investigated how the global perturbation of the brain network using propofol anesthesia alters the spectral slope in individuals in DOC and whether this response contains information about an individual’s pre-anesthetic level of consciousness. Given that the spectral slope in healthy adult EEG steepens in response to propofol anesthesia (Colombo *et al*., 2019; Lendner *et al*., 2020), we hypothesized that larger steepening of the spectral slope in response to anesthesia would be associated with higher levels of consciousness.

We analyzed an existing dataset of 16 individuals in DOC undergoing a protocol of propofol anesthesia (see Methods). We compared 5 minutes of 128-channel EEG recorded prior to the anesthetic protocol (i.e., Baseline state) and during propofol anesthesia (i.e., Anesthesia state). For individuals in an acute DOC (n = 11), recovery of consciousness was assessed three months post-EEG. At this time, five participants had recovered full consciousness (i.e., were able to respond verbally and consistently follow commands), four participants did not recover consciousness and two participants had life-sustaining treatment withdrawn (see Methods). The aperiodic component of the EEG was defined by the offset and slope of the PSD and estimated using the FOOOF algorithm (see Methods). The propofol-induced change of the aperiodic component Δslope and Δoffset was defined by the difference between the value in the Baseline and Anesthesia conditions (Baseline - Anesthesia). In order to capture the aperiodic component of the traditionally assessed bandwidths, we focused on the spectral slope in the 1-45 Hz. Due to prior evidence that the spectral slope in higher frequencies (> 30 Hz) is specifically sensitive on the effect of anesthesia (Colombo *et al*., 2019; Lendner *et al*., 2020), we also assessed the spectral slope in the 30-45 Hz range.

Anesthesia significantly steepened the spectral slope in the 30-45 Hz range (t(15) = 4.554, p < 0.001) and the 1-45 Hz range (t(15) = 2.542, p < 0.05) (see Fig. 2A, 2B). The Δslope in both frequency ranges was dependent on the spectral slope at Baseline, with a flatter spectral slope at Baseline indicating a stronger change in response to propofol (see Fig. S3). Most interestingly, the absolute amount of Δslope (i.e. | Δslope|) in the 30-45 Hz range predicted participant’s CRS-R score (R^2^=0.30, F(1,13)=9.62, p<0.05), with a higher CRS-R score indicating a stronger change of the aperiodic component in response to propofol (see Fig. 2D). Contradicting previous research in healthy adults (Colombo *et al*., 2019; Lendner *et al*., 2020), some participants exhibited a flattening of the spectral slope in response to exposure to anesthesia (see Fig. 2B, see Discussion). However, the correlation to individual’s level of consciousness was also present when considering the directionality of change (R^2^=0.23, F(1,13)=6.33, p<0.05) (i.e. without taking the absolute value) (see Fig, S3). In both cases, participants with a higher CRS-R score exhibited a larger steepening of the spectral slope in response to propofol anesthesia. This effect could not be replicated in the 1-45 Hz range (see. Fig. 2D). While the Δslope in the 30-45 Hz range was homogenously distributed, the slope in the 1-45 Hz range differed between central-parietal and lateral regions (see Fig. 2C). Participants’Δslope did not differ between chronic and acute states (see Fig. S3). All results were replicated using the spectral offset (see Fig. S4).

**Figure 2.**
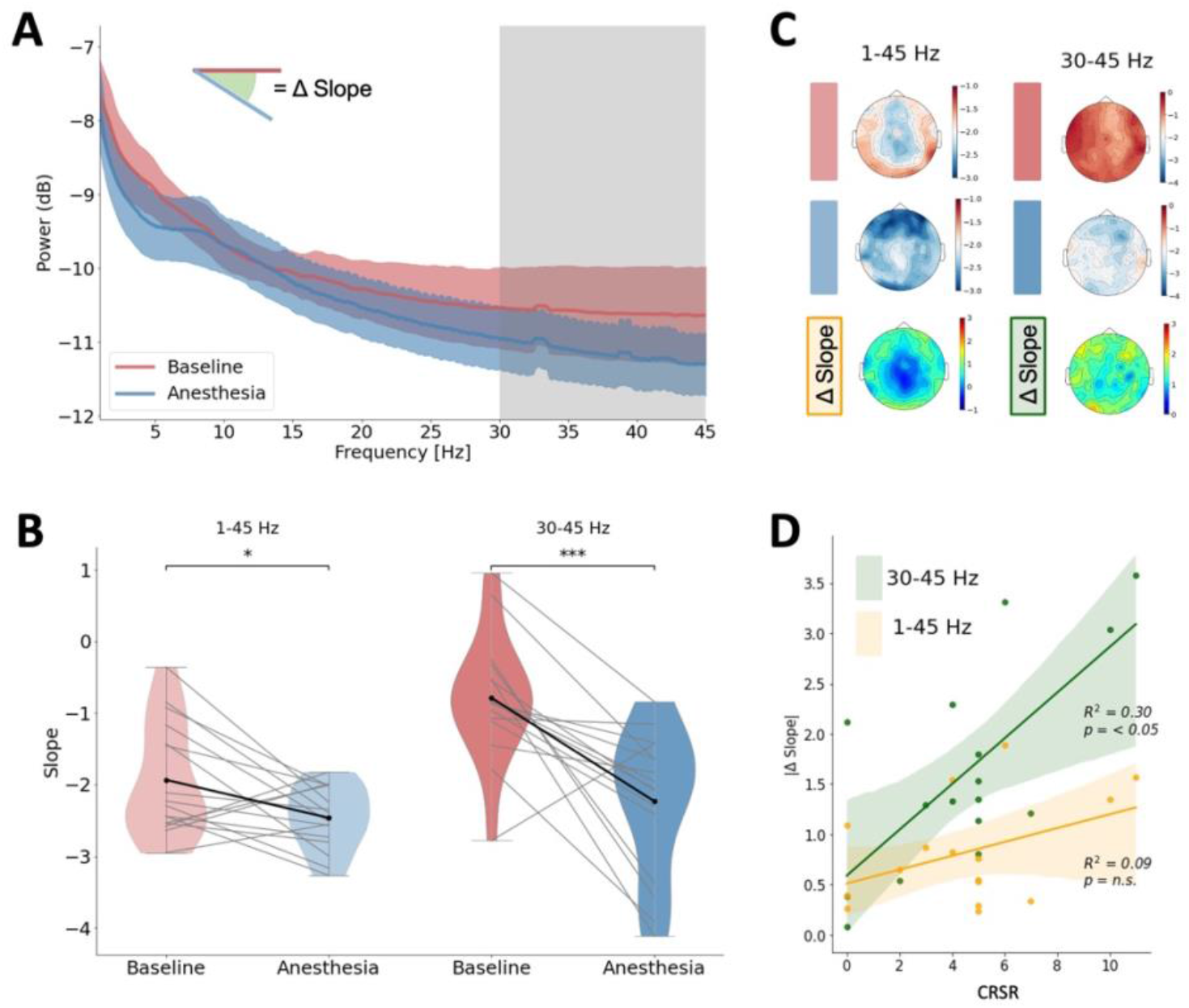
Alterations of the spectral slope in DOC and general anesthesia. **(A)** Power spectral density of Baseline (red) and Anesthesia state (blue), averaged across participants. **(B)** Change in spectral slope in response to propofol anesthesia in 1-45 Hz (left) and 30-45 Hz (right). **(C)** Group-averaged spatial distribution of the spectral slope at Baseline and Anesthesia state and the change in spectral slope (Δ Slope). **(D)** The absolute change in spectral slope over all channels in the 30-45 Hz range correlates with participants’ CRS-R score.

### Propofol-induced loss of chaoticity and information-richness relies on pre-anesthetic EEG spectral slope

To further investigate the mechanism underlying the change in the aperiodic component, we investigated how the spectral slope was related to alterations in network criticality and complexity. Signal complexity was estimated using two variants of Lempel-Ziv complexity (LZC), namely the univariate (Lempel and Ziv, 1976) and concatenated LZC (Schartner *et al*., 2015) (see Methods). To account for the influence of propofol-induced spectral changes on measures of complexity, both variants of LZC were computed with two types of normalization: 1) the classic normalization using randomly shuffled signal, and 2) a phase-normalized signal, which has been previously argued to reduce the bias of LZC to spectral properties of the signal (Toker *et al*., 2022) (see Methods). The network criticality and chaoticity were estimated using the pair correlation function (PCF) (Yoon *et al*., 2015), and the modified 0-1 chaos test (Toker *et al*., 2022), respectively. Based on the signal chaoticity, we additionally estimated the proximity to edge-of-chaos criticality (EOC), following the methods proposed by Toker et al. (2022) (see Methods). The propofol-induced change of the signal complexity and network criticality was defined by the difference between the values in the Baseline and Anesthesia condition (Baseline - Anesthesia).

Anesthesia had no significant group-level effect on any estimate of signal complexity (i.e. concatenated phase-normalized, concatenated shuffle-normalized, univariate phase-normalized, univariate shuffle-normalized) (see Fig. S5). Similarly, no significant group-level effect was observed in signal chaoticity, the PCF, and the closeness to the EOC (see Fig. S5). Instead, individuals in DOC exhibited a heterogenous set of reactions in response to propofol anesthesia (see Fig. S5). Contradicting previous research in healthy adults (Kim and Lee, 2019; Toker *et al*., 2022), some participants exhibited an increase in PCF and complexity, decrease in chaoticity and approach to the EOC under exposure to propofol anesthesia.

We assessed how the anesthetic-induced change of the signal complexity, chaoticity and criticality depended on the brain’s spectral slope at the Baseline state. In the 1-45 Hz range, the spectral slope at Baseline correlated positively with the anesthetic-induced change of the PCF (r=0.53, p<0.05), closeness to EOC (r=0.51, p<0.05), the shuffle-normalized concatenated LZC (r=0.74, p<0.01) and shuffle-normalized univariate LZC (r=0.81, p<0.001) (see Fig.3). The spectral slope in the 1-45 Hz range at Baseline correlated negatively with signal chaoticity (r=-0.57, p<0.05). No correlation was observed with the spectral slope in the 30-45 Hz range. Cumulatively, a flatter spectral slope at Baseline resulted in a stronger decrease of the PCF, stronger distancing form EOC, stronger loss of complexity (i.e. using shuffle-normalized LZC) and stronger increase in chaoticity under exposure to propofol anesthesia. Using the phase-normalization for LZC (i.e. to reduce the bias of spectral changes), the relation to the spectral slope at Baseline neutralized (see Fig. 3), indicating that the shown effects in signal complexity are highly biased by spectral changes of the signal (see Discussion).

**Figure 3.**
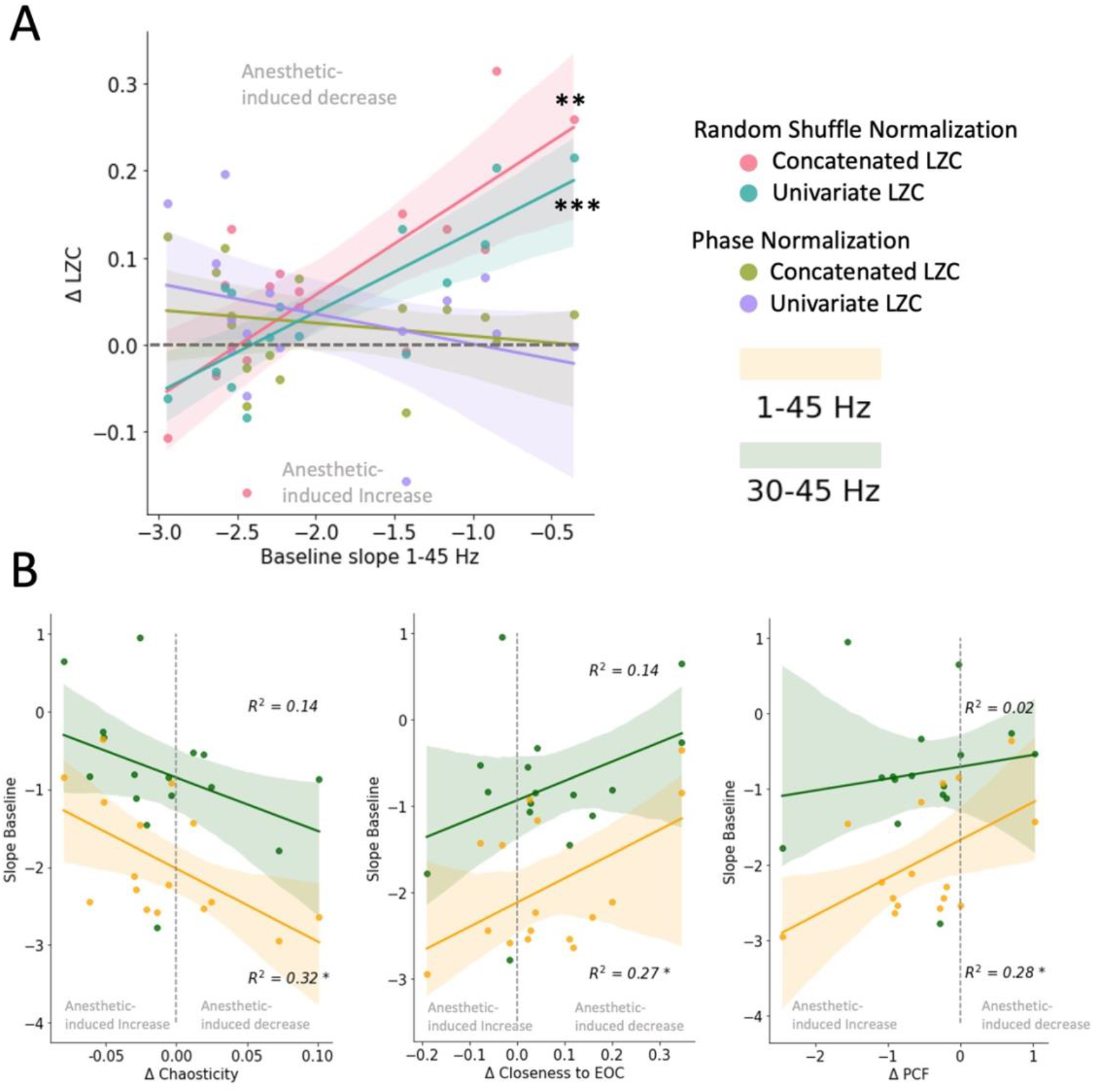
Relation between the spectral slope at baseline and the anesthetic-induced change of **(A)** signal complexity, using four different types of LZC, and ***(B)*** Chaoticity, Closeness to edge of chaos (EOC) and the pair correlation function (PCF). * indicates *p*<0.05, ** indicates *p*<0.01, *** indicates *p*<0.001.

Additionally, we demonstrated how the anesthetic-induced change of the signal complexity, chaoticity and criticality is not only dependent on the properties of the brain at Baseline, but is also reflected in the anesthetic-induced change of the spectral slope. The propofol-induced change of the spectral slope in the 1-45 Hz range correlated positively with the change of the PCF (r=0.56, r<0.05) and negatively with the change of the chaoticity (r=-0.58, r<0.05) (see Fig.S6). No significant correlation was observed between the change of the spectral slope and the change of the closeness to the EOC (see Fig. S6). The change in the spectral slope in the 1-45 Hz range correlated positively with shuffle-normalized concatenated (r=0.56, r<0.05) and univariate LZC (r=0.60, r<0.05), but negatively with phase-normalized concatenated (r=-0.51, r<0.05) and univariate LZC (r=0.57, r<0.05) (see Discussion). Cumulatively, a stronger steepening of the spectral slope in response to propofol anesthesia was accompanied by a stronger loss of signal complexity (i.e. as estimated by shuffle-normalized LZC), criticality (as estimated by decreased PCF) and stronger increase of chaoticity. Surprisingly, the measures of LZC, which were argued to be less biased by the spectral properties of the signal (i.e. using phase-normalization) inversed this relation with a stronger steepening of the spectral slope indicating a stronger information gain (see Discussion).

### The aperiodic EEG component and concatenated complexity during exposure to anesthesia contains prognostic information for individuals in a DOC

During the anesthetized state, the 1-45 Hz aperiodic EEG component distinguished participants who recovered consciousness 3 months post-EEG from those who did not recover consciousness (see Fig. 4A). Participants who recovered consciousness exhibited a significantly steeper spectral slope (t(7) = −3.57, p < 0.01), compared to participants who did not recover consciousness. There was no prognostic value in the spectral slope at Baseline, nor in the propofol-induced change of the aperiodic component (see Fig. 4B). The effect was not replicated in the 30-45 Hz range.

**Figure 4.**
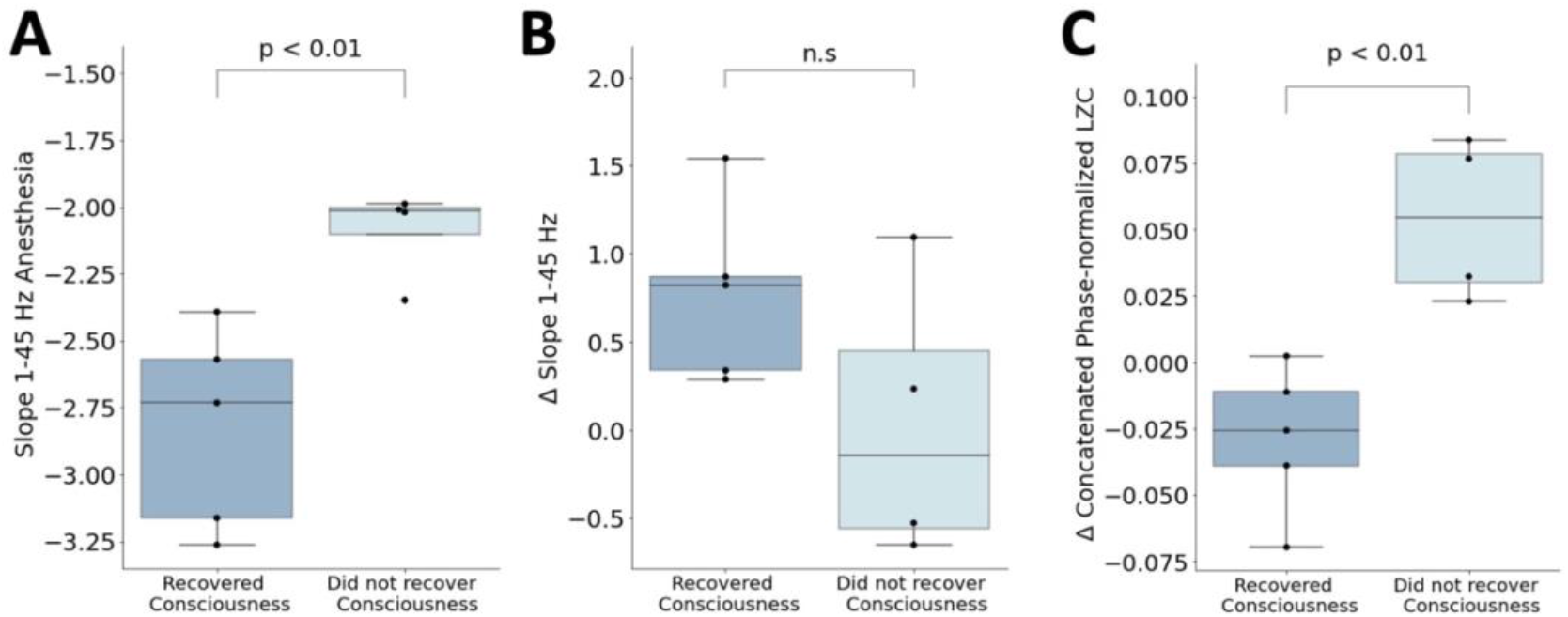
Prognostic value of the spectral slope and signal complexity (A) The spectral slope during anesthesia in the 1-45 Hz range differs between individuals who did and did not recover consciousness 3 months post-EEG. (B) The anesthetic-induced change of the spectral slope does not contain prognostic information for DOC. (C) The anesthetic-induced change of the concatenated LZC contains prognostic information for DOC. Error bars represent standard errors.

Due to the loss of prognostic value in the propofol-induced steepening of the spectral slope, we performed an *ad hoc* analysis investigating the prognostic value in the change of LZC. Most interestingly, this analysis revealed prognostic value in the propofol-induced change of the concatenated LZC (see Fig. 4C). While participants who later recovered consciousness exhibited a propofol-induced increase of phase-normalized concatenated LZC, the signal complexity of participants who did not later recover consciousness decreased in response to anesthesia (t(7) = - 4.18, p < 0.01) (see Discussion). This effect was not replicated using shuffle-normalized LZC. The difference in phase-normalized univariate LZC did not reach significance (t(7) = - 4.23, p=0.06).

## Discussion

This is the first study to investigate the aperiodic component of EEG and its response to propofol for the assessment of level of and capacity for consciousness in individuals in a DOC. We showed that the aperiodic component in the baseline EEG of individuals in a DOC contains diagnostic information above and beyond traditional analysis of periodic EEG components. The significance of this finding is underscored by the fact that although 70% of the DOC participants included in this study did not have an EEG oscillatory peak, analysis of EEG oscillatory power remains by far the most prevalent method for investigating the brain activity in this population. Building upon previous work from our group showing that global brain network perturbation using anesthesia has prognostic value (Duclos *et al*., 2022), we also investigated the diagnostic and prognostic value of the aperiodic EEG component change upon exposure to propofol. We showed that the propofol-induced change of the EEG aperiodic component positively correlated with individual’s pre-anesthetic level of consciousness, with higher levels of consciousness indicating a larger propofol-induced steepening of the spectral slope. We further showed that the pharmacologically induced change of the spectral slope, information-richness and criticality relied on individual’s pre-anesthetic aperiodic component. During exposure to anesthesia, the aperiodic component contained prognostic value for individuals with DOC. Cumulatively, our results highlight the urgent need to reconsider analysis of DOC brain activity in light of the diagnostic and prognostic information contained in the traditionally discarded aperiodic EEG component.

It has been widely proposed that the diagnostic and prognostic power of the EEG aperiodic component should be investigated in brain-injured, unresponsive patients (Colombo *et al*., 2019; Lendner *et al*., 2020). The strong evidence that the aperiodic EEG component distinguishes states of wakefulness from sleep (Lendner *et al*., 2020) and general anesthesia (Colombo *et al*., 2019; Lendner *et al*., 2020) motivated investigations of diagnostic value of this signal for unresponsive patients. In a clinical population, Lanzone et al. (2022) proposed the aperiodic slope as an index for longitudinal recovery from stroke. To date, only a single study by Alnes et al. (2021) assessed the spectral properties of pathologically unresponsive individuals and showed that the aperiodic component of the EEG is altered in patients in a coma, compared to healthy adults. However, EEG in this study was recorded when participants were under varying levels of sedation: this critically affects interpretation of the results, as the non-oscillatory characteristics of DOC are overshadowed by the known effect of anesthesia on the spectral slope. Our study compares the aperiodic EEG component for DOC participants before and during exposure to a targeted and stable concentration of anesthesia, not only dissociating these two potentially confounding factors, but also illustrating the diagnostic potential of the within-subject changes in spectral slope in this population.

Traditionally, DOC has been described through alterations in the power of canonical EEG frequency bands (see Bai, Xia and Li, 2017 as a review). Recent best practice in EEG analysis recommends removing the aperiodic component from the signal prior to power analysis (Donoghue, Schaworonkow and Voytek, 2021). However, the EEG of individuals in DOC is most commonly characterized by a total absence of oscillatory peaks. While this prevents any meaningful oscillation-based analysis, our study demonstrated that the aperiodic component of EEG still contains important information about the individual’s level and capacity for consciousness. However, the results of our study do not support neglecting oscillatory power per se; rather, they highlight the value of the aperiodic component in populations where oscillatory peaks are not systematically present.

Although previous research found significant correlations between participants’ CRS-R score and power in alpha (Chennu *et al*., 2014) and delta bandwidths (Lechinger *et al*., 2013), we did not reproduce these results in the current study. We consider two possible explanations for this discrepancy: first, whereas Chennu et al. (2014) included individuals with a CRS-R score above 7, only 24% of the participants in the present study met this criterion. This suggests that EEG oscillations may increase nonlinearly with an individuals’ level of consciousness, which should be explored in future research. Second, when analyzing power in narrow frequency bands, changes in the power spectrum driven solely by the slope of the aperiodic component can be misinterpreted as a decrease in low frequencies and increase in high frequencies (Donoghue, Schaworonkow and Voytek, 2021). Thus, the previously shown increased delta power in individuals with lower levels of consciousness (Lechinger *et al*., 2013; Chennu *et al*., 2014) might in fact be epiphenomenal to a steepening of the spectral slope, instead of oscillatory power.

Other studies have also presented evidence for the role of EEG oscillatory power in the assessment of DOC. Lechinger et al (2013) demonstrated a correlation between the occipital peak frequency and individuals’ level of consciousness. However, participants who did not exhibit an oscillatory peak were excluded from the analysis (Lechinger *et al*., 2013). We hypothesize that oscillatory power and peak frequency, if present, can play a complementary role to the aperiodic component for the diagnosis of levels of consciousness. Although we cannot test this hypothesis directly in this study, as only 6 out of 43 participants exhibited an oscillatory peak in the 4-13 Hz range, this remains a fruitful area for further research. In the case that no oscillatory peak is present, our results demonstrate that the diagnostic value of EEG for individuals in DOC might be fully attributable to the aperiodic component.

Despite strong evidence that the spectral slope is related to consciousness (Colombo *et al*., 2019; Lendner *et al*., 2020), there is disagreement about whether it is a measure of arousal (i.e. vigilance) (Lendner *et al*., 2020) or of consciousness level (i.e. awareness) (Colombo *et al*., 2019). Rather than framing our results within these dimensions – which have been criticized as failing to represent the multifaceted nature of consciousness and its disorders (Bayne, Hohwy and Owen, 2016) — we focus on a mechanistic interpretation of the aperiodic component and its link to consciousness. Using *in silico* modeling, the slope of the aperiodic component has been suggested to be a marker of the network’s E/I balance (Gao, Peterson and Voytek, 2017; Medel *et al*., 2020). Moreover, consciousness has been proposed to be underpinned by an optimal E/I balance (Toker *et al*., 2022), which tunes the brain towards a state of criticality and information-richness (Shew *et al*., 2011). When this balance is disrupted (i.e. more inhibition than excitation) the network diverges from criticality, exhibits a steeper spectral slope (Gao, Peterson and Voytek, 2017; Medel *et al*., 2020; see Zimmern, 2020 as a review) and reduced signal complexity (Medel *et al*., 2020). Our results further support the proposition that the brain of individuals in DOC operates far from a critical point (Toker *et al*., 2022), resulting in weaker network susceptibility to global perturbations, such as propofol anesthesia.

Exposure to the inhibitory drug propofol causes steepening of the spectral slope (Colombo *et al*.,2019; Lendner *et al*., 2020), reflecting the brain’s shift towards inhibition. In this study, the anesthetic-induced change of the spectral slope correlated with participants’ CRS-R score and thus, depended on their baseline level of consciousness. One potential explanation for this phenomenon is that the effect of propofol depends on the pre-anesthetic E/I balance. While exposure to propofol has a large effect on a well-balanced brain, an imbalanced brain may have a reduced capacity to shift, as it is closer to maximum imbalance. In other words, the already-steepened spectral slope in comatose patients could reflect the brain’s high imbalance and distance from criticality, resulting in a reduced response to propofol anesthesia. Comparably, stronger brain network reconfiguration in the alpha bandwidth following exposure to propofol has been linked to higher potential for recovery of consciousness (Duclos *et al*., 2022). Thus, one potential interpretation of our results is that individuals in DOC are characterized by an E/I imbalance, with higher imbalance resulting in a reduced reaction of the aperiodic component to general anesthesia and lower levels of consciousness.

Despite the group-level steepening of the spectral slope during exposure to anesthesia, some individuals in DOC exhibited an alternative pattern: a flatter spectral slope in the anesthesia state, accompanied by a more complex signal and increased criticality. Toker et al. (2022) observed a similar inconsistency, with one individual in DOC exhibiting increased chaoticity after regaining consciousness. Although an increase of complexity and flattening of the spectral slope following exposure to anesthesia is counterintuitive for healthy individuals, DOC might be a heterogenous set of conditions, which might be either too chaotic or too stable. The spectral slope in the 1-45 Hz range at baseline clearly separated individuals with DOC into two groups: individuals with flatter slopes exhibited a propofol-induced steepening of the slope, while individuals with slopes steeper than 2.6 showed a countertrend flattening of the spectral slope (see Fig. 2B). Further characterization of the clinical differences between these groups is warranted in future research.

Pharmacologically-induced reduction of signal complexity has been previously demonstrated in numerous studies on healthy individuals using propofol, isoflurane and sevoflurane (Zhang, Roy and Jensen, 2001; Schartner *et al*., 2015; Hudetz *et al*., 2016; Wang *et al*., 2017; Toker *et al*., 2021). However, in this study, we could not identify a group-level decrease of LZC. In contrast, we demonstrated that the loss of signal complexity was strongly related to the signal’s spectral properties prior to exposure to anesthesia and in response to anesthesia. Whereas the relation between the spectral slope and signal complexity has been previously shown using modeling studies (Medel *et al*., 2020), this study provides the first evidence from a clinical population. Most interestingly, the relation between the spectral slope and the signal complexity vanished or inversed after normalizing the LZC with phase-randomized surrogate, which was argued to be most robust to spectral changes of the signal (Toker *et al*., 2022). This suggests that observed changes of LZC - which are commonly normalized using signal length or a randomly shuffled signal - might be largely driven by spectral changes after exposure to anesthesia. An independent investigation of anesthetic-induced loss of signal complexity and change of the aperiodic component for the evaluation of levels of consciousness would be recommended for future research.

The results of this study should be interpreted in light of several limitations. First, participants were assessed using the JFK Coma Recovery Scale-Revised (CRS-R) (Kalmar and Giacino, 2005), which is a behavioral scale for the assessment of responsiveness. The resulting score is widely used as a surrogate measure of consciousness. However in DOC, consciousness can be fully dissociated from behavior (Owen *et al*., 2006; Sanders *et al*., 2012; Mashour and Avidan, 2013). Thus, it is possible that the true level of consciousness in study participants was not accurately captured by this metric and that true levels of consciousness were underestimated by the behavioral score of responsiveness. Whereas the first part of this study (Baseline recording) cannot differentiate whether the aperiodic component captures levels of consciousness or responsiveness, the value of the aperiodic component for detecting covert consciousness in unresponsive individuals could be explored in future research. In the second part of this study, DOC participants underwent a protocol of propofol anesthesia. Independent of the pre-anesthetic level of responsiveness, the individual’s anesthetic-induced brain reaction and changes in the spectral slope are attributable to loss of consciousness, which might diverge from the degree of anesthetic-induced loss of responsiveness. Thus, a strong brain reaction to anesthetics despite low to no behavioral changes, might be an indicator for higher pre-anesthetic levels of consciousness reaching beyond the estimated level of responsiveness. This possibility should be more closely investigated in future research.

Second, participants in this study were recruited after a variety of brain injuries, including stroke, traumatic and anoxic brain injury. Although the spectral slope is steeper in the hemisphere most affected by a stroke, (Lanzone *et al*., 2022), this study did not account for the location of the brain injury. Additionally, the trend towards low CRS-R score within this study’s participants led to high imbalance between the classical diagnostic groups (i.e., coma, unresponsive wakefulness syndrome, minimally conscious state, emergence). We therefore did not perform a group-comparative statistical analysis to distinguish the classical diagnostic categories. Third, this study explored the link between the aperiodic component and network criticality, measured by the PCF (Kim and Lee, 2019) and the modified 01-chaos test (Toker *et al*., 2022). The PCF specifically in the alpha frequency range has shown a strong link to measures of consciousness (Lee *et al*., 2018; Kim and Lee, 2019). Despite the lack of an alpha peak the PCF in this study was estimated in the 8-13 Hz range. An influence of increased alpha-power during exposure to propofol on the estimate of PCF cannot be excluded. Following the provided methods by Toker et al. (2022) the modified 0-1 chaos test was only performed on channels which exhibited an oscillatory peak in the 1-6 Hz range. There is a large variety of measures within the methodological framework of network criticality (see Zimmern, 2020 as a review); the broader exploration of the aperiodic component and its relation to different measures of criticality, which do not rely on oscillation, is strongly recommended for future research. For the estimation of the closeness to EOC, we used an alpha of 0.85, as proposed by Toker et al. (2022). This value has been empirically defined to distinguish levels of wakefulness from unconscious states. A validation or refinement of this value in a larger clinical population would be recommended in further research. Fourth, the detection of oscillatory peaks in the first part of the analysis was performed on the electrode-averaged spectrum. Whereas most individual’s electrode-averaged PSD did not exhibit oscillatory peaks, we cannot exclude the possibility of small oscillatory peaks on the level of single electrodes.

In conclusion, we have demonstrated the value of the aperiodic EEG component for the assessment of individuals in a DOC following brain injury. At Baseline, individuals with a lower level of consciousness exhibit a steeper spectral slope (i.e., faster decay of power over frequency). The anesthetic-induced change in the aperiodic component depends on individual’s pre-anesthetic level of consciousness and accompanies the brain’s loss of criticality. The aperiodic EEG component has been historically discarded; this research highlights a critical need to reconsider the traditional treatment of this component of the EEG in research with individuals in DOC.

## Materials and Methods

### Participants and anesthetic protocol

This study combined two existing datasets of DOC participants (i.e., one dataset of Baseline EEG recordings and one dataset of individuals in DOC, undergoing an anesthetic protocol). In total, 43 individuals in a DOC (22 male, 42 ±15.13 years old) were included in this study. Individuals in a DOC were included following acquired brain injury (anoxic, traumatic, hypoxic brain injury, stroke) and assessed by a trained experimenter using the Coma Recovery Scale-Revised (CRS-R) (Kalmar and Giacino, 2005). Participants were excluded if they were receiving sedation at the time of the study. For all participants, written informed consent was provided by their legal representative in accordance with the Declaration of Helsinki. The study was approved by the McGill University Health Center Research Ethics Board (15-996-MP-CUSM) and the Western University Health Science Research Ethics Board (Project ID 100628). Among the DOC participants, 14 were in MCS, 25 in UWS and four in a coma (CRS-R = 5.88 ± 4.02). For participants in an acute DOC (n = 18), clinical outcomes were assessed three months post recording. At this time, six participants had recovered full consciousness (i.e., were able to respond verbally and consistently follow commands). Nine participants did not recover consciousness remained in a DOC. Three participants had life-sustaining treatment withdrawn and were not excluded from the analysis of prognostic value.

Sixteen (n=16) of the above described individuals in a DOC (5 male, 44 ±18.24 years old) underwent an anesthetic protocol, explained in (Blain-Moraes *et al*., 2016). Briefly, participants were anesthetised with propofol at a target effect site concentration of 2.0 μg/ml. In this study, we include a period of 5 minutes resting state prior to the start of the anesthetic protocol (referred to as: Baseline state) as well as the period of 5 minutes during the infusion of propofol, after the effect site concentration of 2.0 μg/ml has been reached (referred to as: Anesthetized state). Within this subset 11 individuals were in an acute state. Within three months post recording, five participants had recovered full consciousness (i.e., were able to respond verbally and consistently follow commands), four participants did not recover consciousness and two participants had life-sustaining treatment withdrawn.

### Electroencephalography data

Data in both datasets were recorded from a 128-channel EGI Sensor Net using an Amps 400 amplifier (Electrical Geodesic, Inc., USA), a sampling rate of 1 kHz and vertex reference. Electrode impedance was reduced to below 5KΩ prior to data collection. Two participants were recorded using a 64-channel EEG system. Prior to analysis, the raw signal was filtered between 0.5 to 55 Hz, average referenced and resampled to 250 Hz. A notch filter was applied at 60 Hz. Channels with an excessive level of noise were removed prior to average referencing. Non-brain channels were removed from the subsequent analysis. The signal was then epoched in non-overlapping segments of 10 seconds. The signal was visually inspected by a trained investigator to manually reject epochs containing nonphysiologically artifacts. All preprocessing steps were performed using the MNE python toolbox (Gramfort *et al*., 2013).

### Spectral slope and power analysis

The power spectra were calculated for every electrode and epoch, using the Multitaper approach (Prerau *et al*., 2017). All spectral estimates were performed using a frequency range from 0.5 to 50 Hz and a frequency smoothing of ±0.5 Hz, resulting in the use of 9 discrete prolate slepian sequences (dpss) tapers (Prerau *et al*., 2017). The aperiodic component was defined by the EEG spectral slope and offset which were calculated using the FOOOF package (Donoghue *et al*., 2020). This algorithm parametrizes the EEG into an aperiodic and oscillatory component. The aperiodic component of the PSD over frequencies F is defined by:

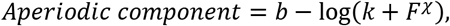

where b is the spectral offset, k is the knee parameter and *χ* is the spectral slope. To be coherent with previous research (Colombo *et al*., 2019; Lendner *et al*., 2020), the spectral slope was herein defined by -*χ*. In the 1-45 Hz range, the algorithm was fit using the fixed and knee aperiodic mode (min_peak_height=0.1, max_n_peaks=10). Whereas the knee mode fits an additional parameter k to the data, this parameter is set to 0 in the fixed mode. In the 30-45 Hz range we exclusively used the fixed mode to fit the data. This can be justified by the narrower frequency band which does not require a knee fit. The results from the model using the knee mode yield an overall better model fit (see Fig. S1) and was presented in this paper. The model error and analysis using the knee mode is provided in the supplementary material (see Fig. S1). The power spectra, as well as the spectral slope were calculated for every participant, epoch, and electrode independently and averaged subsequently. Is it to note that estimating the aperiodic component on the time-averaged PSD did not affect any of the results.

Power analysis in the delta (1-4 Hz), theta (4-8 Hz), alpha (8-13 Hz), beta (13-30 Hz) and gamma (30-45 Hz) bands was performed before and after the removal of the broadband aperiodic component from the PSD. In the first part, oscillatory power was calculated using the relative contribution of one frequency band to the overall PSD. In the second step, the PSD was flattened (i.e. removal of the aperiodic component) using the FOOOF package (Donoghue *et al*., 2020). The resulting oscillatory power was averaged within each frequency band. To identify whether the PSD of a signal participant exhibits a peak frequency, a separate model was fit on the electrode-averaged PSD. The propofol-induced change of the spectral slope was defined as the distance between the baseline value and the value in the anesthetized state (i.e. Baseline - Anesthesia).

### Complexity and Criticality

Signal information-richness was calculated using four measures of LZC. All estimates of LZC were calculated on every epoch individually and averaged subsequently. For all estimates of LZC, the signal was low-pass filtered at 45 Hz and binarized using the mean of its instantaneous amplitude. For the univariate LZC, complexity was calculated on every channel individually and averaged subsequently, using the median over all channels in one epoch (Lempel and Ziv, 1976). For the concatenated LZC, the signal of all channels in one epoch was concatenated before compression, as described by Schartner et al. (2015).

To account for biases of the spectral properties on the signal complexity, both measures of LZC were normalized using two approaches 1) shuffle-normalization and 2) phase-normalization. In the first approach the LZC of every epoch is normalized by the complexity of the randomly shuffled binarized time-series (Schartner *et al*., 2015). The second method compares the signal’s LZC to a phase-randomized surrogate of this signal and has been demonstrated to be most robust against spectral changes of the signal (Toker *et al*., 2022). Both estimates of LZC were calculated using the code provided by Toker et al (2022) and an additional custom function for shuffle-normalization.

Criticality of the brain network was defined using the pair correlation function (PCF), which is an estimate of global phase synchronization or network susceptibility (Yoon *et al*., 2015; Lee *et al*., 2018; Kim and Lee, 2019). Due to its previously demonstrated link to levels of consciousness and the information integration of the underlying brain network (Lee *et al*., 2018; Kim and Lee, 2019), we estimated the PCF in the 8-13 Hz range. The PCF was estimated for every epoch individually and averaged subsequently.

To provide a second measure of network criticality, we further estimated the network’s closeness to the edge of chaos. Chaoticity was estimated using the modified 0-1 chaos test (Gottwald and Melbourne, 2004; Toker *et al*., 2022). Following the recommendations of Toker et al. (Toker *et al*.,2022), signal chaoticity was estimated on the low-frequency cortical activity. We therefore applied the FOOOF algorithm on every channel and epoch individually to identify the highest peak frequency between 1 to 6 Hz. Channels without an oscillatory peak in this frequency range were removed from chaoticity analysis (this was the case for 22.90 % of the signal, see Discussion). The edge of chaos criticality was estimated using the methods provided by Toker et al. (2022) and the proposed alpha of 0.85 (see Discussion). The propofol-induced change in both measures was defined as the difference between the baseline value to the value in the anesthetized state.

### Statistical analysis

Statistical analysis was performed using Python. In the first section, the link between the oscillatory and aperiodic component with participants’ CRS-R score was assessed using a multivariate linear regression, preconditioned on participants age. Assumptions were tested using Shapiro-Wilk test. Reported R^2^ values correspond to the adjusted R^2^. For visualization only, individual R^2^ were obtained using univariate linear regression. Confidence intervals were estimated using bootstrapping (1000 iterations). In the second part, the change of the spectral slope, criticality, complexity and chaoticity in response to propofol anesthesia was assessed using repeated measures t-test. P-values were corrected using Bonferroni-correction. The relation between Δslope and individual’s CRS-R score was estimated using Linear Regression, preconditioned on participants age. To avoid effects of multicollinearity the model was fit on the 1-45 Hz and 30-45 Hz range individually. The effect between the spectral slope at baseline and its propofol-induced change was assessed using Pearson correlation. Group differences between recovered and non-recovered participants were assessed using independent t-test. All p-values were corrected using Bonferroni-correction.

## Acknowledgments

The authors thank all participants for their involvement in the study. They also wish to thank Uncheol Lee for providing code for the criticality analysis, Caroline Arbour for her assessments and training on the CRS-R, as well as Yacine Mahdid, Danielle Nadin, Alexander Rokos, Jason da Silva Castanheira, Laura Gonzalez-Lara, Allison Frantz and Miriam Han for their involvement in participant recruitment and data acquisition. AMO is a CIFAR Fellow. SBM is supported by the Canada Research Chairs Program (Tier II). CD is a Junior 1 Research Scholar from the Fonds de recherche du Québec – Santé (FRQS). This research is supported in part by the FRQNT Strategic Clusters Program (2020-RS4-265502 - Centre UNIQUE - Union Neurosciences & Artificial Intelligence – Quebec. This study was funded through an NSERC Discovery Grant (RGPIN-201603817), the Canada Excellence Research Chairs Program (#215063), the Canadian Institutes of Health Research (#408004). This research was undertaken thanks in part to funding from the Canada First Research Excellence Fund and Fonds de recherche du Québec, awarded to the Healthy Brains, Healthy Lives initiative at McGill University, and the International Anesthesia Research Society (IARS).

## References

Alnes, S.L. et al. (2021) ‘Complementary roles of neural synchrony and complexity for indexing consciousness and chances of surviving in acute coma’, NeuroImage, 245, p. 118638. Available at: https://doi.org/10.1016/j.neuroimage.2021.118638.

Bai, Y., Xia, X. and Li, X. (2017) ‘A Review of Resting-State Electroencephalography Analysis in Disorders of Consciousness’, Frontiers in Neurology, 8. Available at: https://doi.org/10.3389/fneur.2017.00471.

Bayne, T., Hohwy, J. and Owen, A.M. (2016) ‘Are There Levels of Consciousness?’, Trends in Cognitive Sciences, 20(6), pp. 405–413. Available at: https://doi.org/10.1016/j.tics.2016.03.009.

Beggs, J.M. and Plenz, D. (2003) ‘Neuronal Avalanches in Neocortical Circuits’, Journal of Neuroscience, 23(35), pp. 11167–11177. Available at: https://doi.org/10.1523/JNEUROSCI.23-35-11167.2003.

Blain-Moraes, S. et al. (2016) ‘Normal Brain Response to Propofol in Advance of Recovery from Unresponsive Wakefulness Syndrome’, Frontiers in Human Neuroscience, 10. Available at: https://doi.org/10.3389/fnhum.2016.00248.

Carhart-Harris, R.L. et al. (2014) ‘The entropic brain: a theory of conscious states informed by neuroimaging research with psychedelic drugs’, Frontiers in Human Neuroscience, 8. Available at: https://doi.org/10.3389/fnhum.2014.00020.

Carhart-Harris, R.L. (2018) ‘The entropic brain - revisited’, Neuropharmacology, 142, pp. 167–178. Available at: https://doi.org/10.1016/j.neuropharm.2018.03.010.

Casali, A.G. et al. (2013) ‘A Theoretically Based Index of Consciousness Independent of Sensory Processing and Behavior’, Science Translational Medicine, 5(198), pp. 198ra105–198ra105. Available at: https://doi.org/10.1126/scitranslmed.3006294.

Chennu, S. et al. (2014) ‘Spectral Signatures of Reorganised Brain Networks in Disorders of Consciousness’, PLOS Computational Biology, 10(10), p. e1003887. Available at: https://doi.org/10.1371/journal.pcbi.1003887.

Colombo, M.A. et al. (2019) ‘The spectral exponent of the resting EEG indexes the presence of consciousness during unresponsiveness induced by propofol, xenon, and ketamine’, NeuroImage, 189, pp. 631–644. Available at: https://doi.org/10.1016/j.neuroimage.2019.01.024.

Donoghue, T. et al. (2020) ‘Parameterizing neural power spectra into periodic and aperiodic components’, Nature Neuroscience, 23(12), pp. 1655–1665. Available at: https://doi.org/10.1038/s41593-020-00744-x.

Donoghue, T., Schaworonkow, N. and Voytek, B. (2021) ‘Methodological considerations for studying neural oscillations’, European Journal of Neuroscience, pp. 1–26. Available at: https://doi.org/10.1111/ejn.15361.

Duclos, C. et al. (2022) ‘Brain Responses to Propofol in Advance of Recovery from Coma and Disorders of Consciousness: A Preliminary Study’, American Journal of Respiratory and Critical Care Medicine, 205(2), pp. 171–182. Available at: https://doi.org/10.1164/rccm.202105-1223OC.

Gao, R., Peterson, E.J. and Voytek, B. (2017) ‘Inferring synaptic excitation/inhibition balance from field potentials’, NeuroImage, 158, pp. 70–78. Available at: https://doi.org/10.1016/j.neuroimage.2017.06.078.

Gottwald, G.A. and Melbourne, I. (2004) ‘A new test for chaos in deterministic systems’, Proceedings of the Royal Society of London. Series A: Mathematical, Physical and Engineering Sciences, 460(2042), pp. 603–611. Available at: https://doi.org/10.1098/rspa.2003.1183.

Gramfort, A. et al. (2013) ‘MEG and EEG data analysis with MNE-Python’, Frontiers in Neuroscience, 7. Available at: https://www.frontiersin.org/article/10.3389/fnins.2013.00267(Accessed: 23 March 2022).

Hudetz, A.G. et al. (2016) ‘Propofol anesthesia reduces Lempel-Ziv complexity of spontaneous brain activity in rats’, Neuroscience Letters, 628, pp. 132–135. Available at: https://doi.org/10.1016/j.neulet.2016.06.017.

Kalmar, K. and Giacino, J.T. (2005) ‘The JFK coma recovery scale—revised’, Neuropsychological Rehabilitation, 15(3–4), pp. 454–460. Available at:https://doi.org/10.1080/09602010443000425.

Kim, H. and Lee, U. (2019) ‘Criticality as a Determinant of Integrated Information Φ in Human Brain Networks’, Entropy, 21(10), p. 981. Available at: https://doi.org/10.3390/e21100981.

Lanzone, J. et al. (2022) ‘EEG spectral exponent as a synthetic index for the longitudinal assessment of stroke recovery’, Clinical Neurophysiology [Preprint]. Available at: https://doi.org/10.1016/j.clinph.2022.02.022.

Lechinger, J. et al. (2013) ‘CRS-R score in disorders of consciousness is strongly related to spectral EEG at rest’, Journal of Neurology, 260(9), pp. 2348–2356. Available at: https://doi.org/10.1007/s00415-013-6982-3.

Lee, U. et al. (2018) ‘Functional Brain Network Mechanism of Hypersensitivity in Chronic Pain’, Scientific Reports, 8(1), p. 243. Available at: https://doi.org/10.1038/s41598-017-18657-4.

Lempel, A. and Ziv, J. (1976) ‘On the Complexity of Finite Sequences’, IEEE Transactions on Information Theory, 22(1), pp. 75–81. Available at: https://doi.org/10.1109/TIT.1976.1055501.

Lendner, J.D. et al. (2020) ‘An electrophysiological marker of arousal level in humans’, eLife, 9, p. e55092. Available at: https://doi.org/10.7554/eLife.55092.

Mashour, G.A. and Avidan, M.S. (2013) ‘Capturing covert consciousness’, The Lancet, 381(9863), pp. 271–272. Available at: https://doi.org/10.1016/S0140-6736(13)60094-X.

Medel, V. et al. (2020) Complexity and 1/f slope jointly reflect cortical states across different E/I balances, p. 2020.09.15.298497. Available at: https://doi.org/10.1101/2020.09.15.298497.

Muthukumaraswamy, S.D. and Liley, D.TJ. (2018) ‘1/f electrophysiological spectra in resting and drug-induced states can be explained by the dynamics of multiple oscillatory relaxation processes’, NeuroImage, 179, pp. 582–595. Available at: https://doi.org/10.1016/j.neuroimage.2018.06.068.

O’Byrne, J. and Jerbi, K. (2022) ‘How critical is brain criticality?’, Trends in Neurosciences[Preprint]. Available at: https://doi.org/10.1016/j.tins.2022.08.007.

Owen, A.M. et al. (2006) ‘Detecting Awareness in the Vegetative State’, Science, 313(5792), pp. 1402–1402. Available at: https://doi.org/10.1126/science.1130197.

Prerau, M.J. et al. (2017) ‘Sleep Neurophysiological Dynamics Through the Lens of Multitaper Spectral Analysis’, Physiology, 32(1), pp. 60–92. Available at: https://doi.org/10.1152/physiol.00062.2015.

Purdon, P.L. et al. (2013) ‘Electroencephalogram signatures of loss and recovery of consciousness from propofol’, Proceedings of the National Academy of Sciences, 110(12), pp. E1142–E1151. Available at: https://doi.org/10.1073/pnas.1221180110.

Sanders, R.D. et al. (2012) ‘Unresponsiveness ≠ Unconsciousness’, Anesthesiology, 116(4), pp. 946–959. Available at: https://doi.org/10.1097/ALN.0b013e318249d0a7.

Schartner, M. et al. (2015) ‘Complexity of Multi-Dimensional Spontaneous EEG Decreases during Propofol Induced General Anaesthesia’, PLOS ONE, 10(8), p. e0133532. Available at: https://doi.org/10.1371/journal.pone.0133532.

Shew, W.L. et al. (2011) ‘Information Capacity and Transmission Are Maximized in Balanced Cortical Networks with Neuronal Avalanches’, Journal of Neuroscience, 31(1), pp. 55–63. Available at: https://doi.org/10.1523/JNEUROSCI.4637-10.2011.

Swisher, C.B. and Sinha, S.R. (2016) ‘Utilization of Quantitative EEG Trends for Critical Care Continuous EEG Monitoring: A Survey of Neurophysiologists’, Journal of Clinical Neurophysiology, 33(6), pp. 538–544. Available at: https://doi.org/10.1097/WNP.0000000000000287.

Timmermann, C. et al. (2019) ‘Neural correlates of the DMT experience assessed with multivariate EEG’, Scientific Reports, 9(1), p. 16324. Available at: https://doi.org/10.1038/s41598-019-51974-4.

Toker, D. et al. (2021) ‘Consciousness is supported by near-critical cortical electrodynamics’, bioRxiv, p. 2021.06.10.447959. Available at: https://doi.org/10.1101/2021.06.10.447959.

Toker, D. et al. (2022) ‘Consciousness is supported by near-critical slow cortical electrodynamics’, Proceedings of the National Academy of Sciences, 119(7). Available at: https://doi.org/10.1073/pnas.2024455119.

Voytek, B. et al. (2015) ‘Age-Related Changes in 1/f Neural Electrophysiological Noise’, Journal of Neuroscience, 35(38), pp. 13257–13265. Available at: https://doi.org/10.1523/JNEUROSCI.2332-14.2015.

Wang, J. et al. (2017) ‘Suppressed neural complexity during ketamine- and propofol-induced unconsciousness’, Neuroscience Letters, 653, pp. 320–325. Available at: https://doi.org/10.1016/j.neulet.2017.05.045.

Yoon, S. et al. (2015) ‘Critical behavior of the relaxation rate, the susceptibility, and a pair correlation function in the Kuramoto model on scale-free networks’, Physical Review. E, Statistical, Nonlinear, and Soft Matter Physics, 91(3), p. 032814. Available at: https://doi.org/10.1103/PhysRevE.91.032814.

Zhang, X.-S., Roy, R.J. and Jensen, E.W. (2001) ‘EEG complexity as a measure of depth of anesthesia for patients’, IEEE Transactions on Biomedical Engineering, 48(12), pp. 1424–1433. Available at: https://doi.org/10.1109/10.966601.

Zimmern, V. (2020) ‘Why Brain Criticality Is Clinically Relevant: A Scoping Review’, Frontiers in Neural Circuits, 14. Available at: https://www.frontiersin.org/article/10.3389/fncir.2020.00054 (Accessed: 3 March 2022).

