## supplementary material for "Aperiodic brain activity and response to anesthesia vary in disorders of consciousness"

This PDF file includes:

Figures S1 to S6

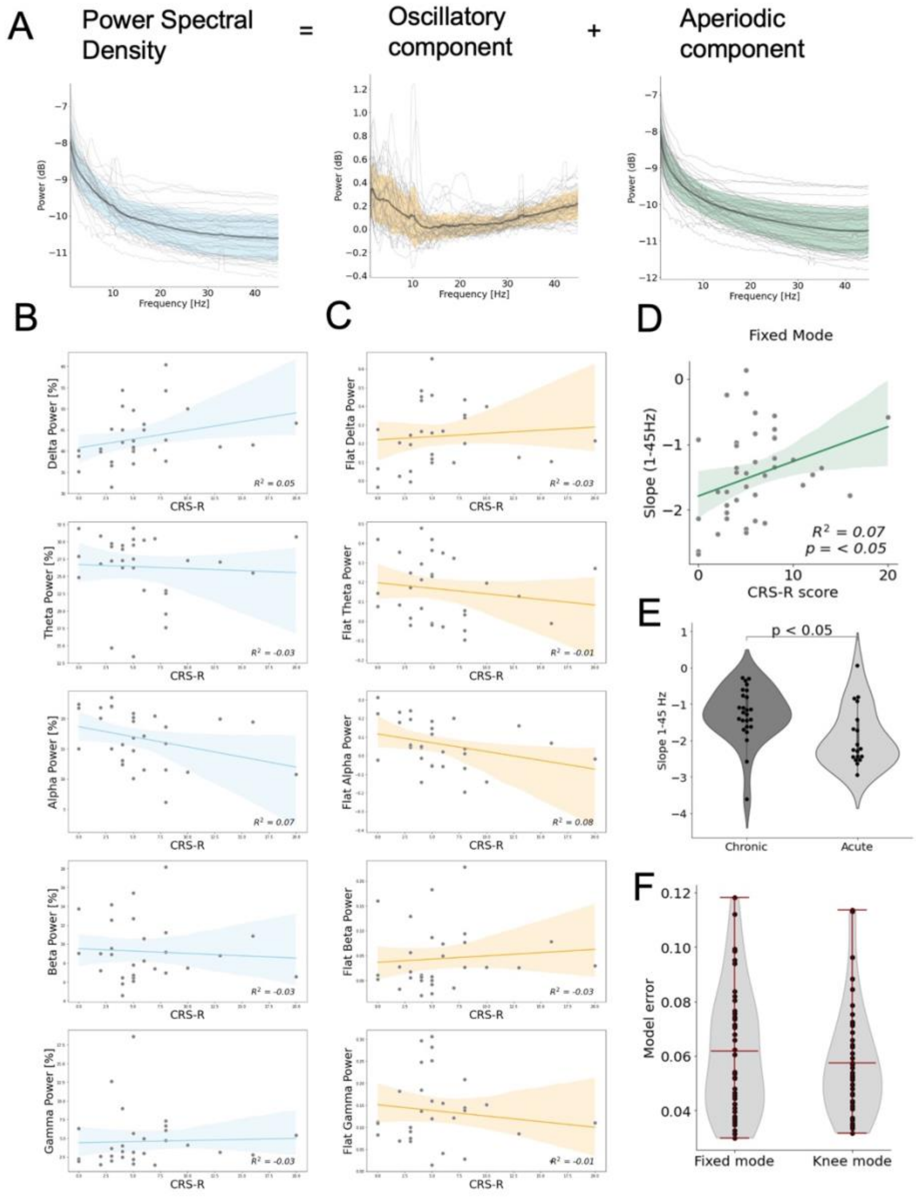


**Fig. S1.** Diagnostic value of EEG spectral properties in DOC. **(A)** The power spectral density of individuals in DOC (blue) separates into an oscillatory component (orange) and an aperiodic component (green). Light grey lines represent individual subjects; darker lines represent the group average and standard deviation. **(B)** The traditional power analysis of oscillatory EEG in the alpha bandwidth does not predict individual's level of responsiveness, as measured by the CRS-R score. **(C)** After removal of the aperiodic component, remaining oscillatory power of the EEG in the alpha bandwidth does not predict individuals' level of responsiveness, as measured by the CRS-R score. **(D)** Replication of the aperiodic component's diagnostic value using the 'fixed mode' instead of 'knee mode'. **(E)** Spectral slope for individuals in acute and chronic DOC. **(F)** Reduced model fitting error when using the model in the 'knee mode'.

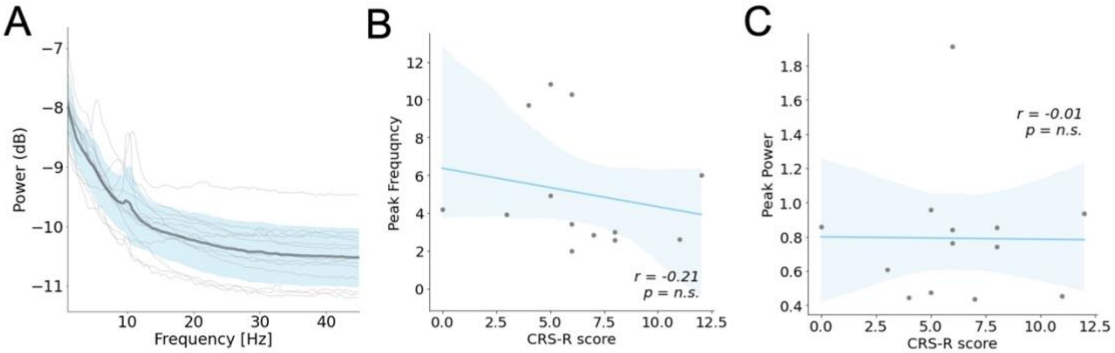


**Fig. S2.** Characteristic of the identified oscillatory peak in 13 individuals. **(A)** The power spectral density of 13 individuals in DOC, light grey lines represent individual subjects, dark grey line represents the group average. **(B)** The identified peak frequency does not correlate with participants CRS-R score. **(C)** The power of the identified peak does not correlate with participants CRS-R score.

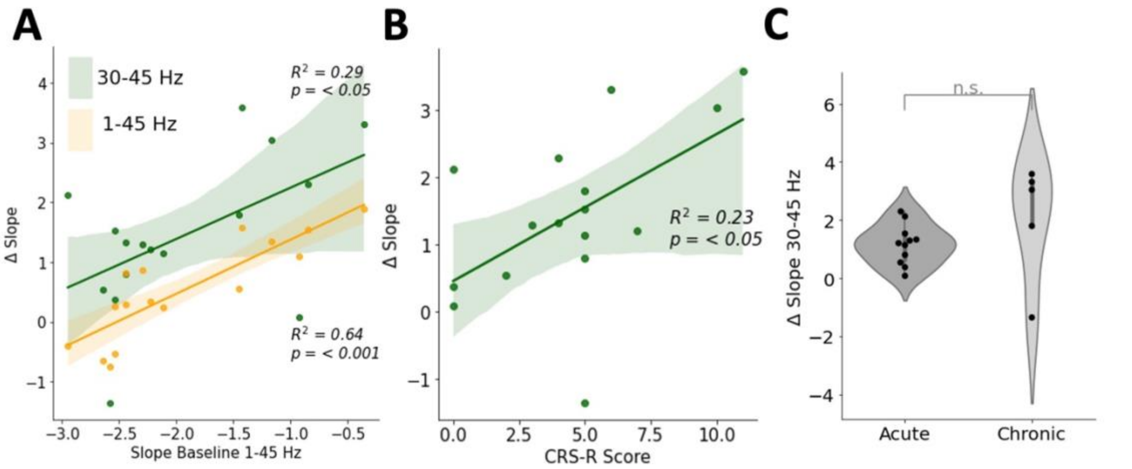


**Fig. S3.** Alterations of the spectral slope under general anesthesia in individuals with disorder of consciousness. **(A)** Flatter spectral slope in the 1-45 Hz range at baseline results in larger anesthetic-induced change of the spectral slope (Δslope) in the 1-45 Hz (yellow) and 30-45 Hz range (green). **(B)** Change in spectral slope in the 30-45 Hz range correlates with individual's level of consciousness. **(C)** Patients Δslope did not differ between chronic and acute states.

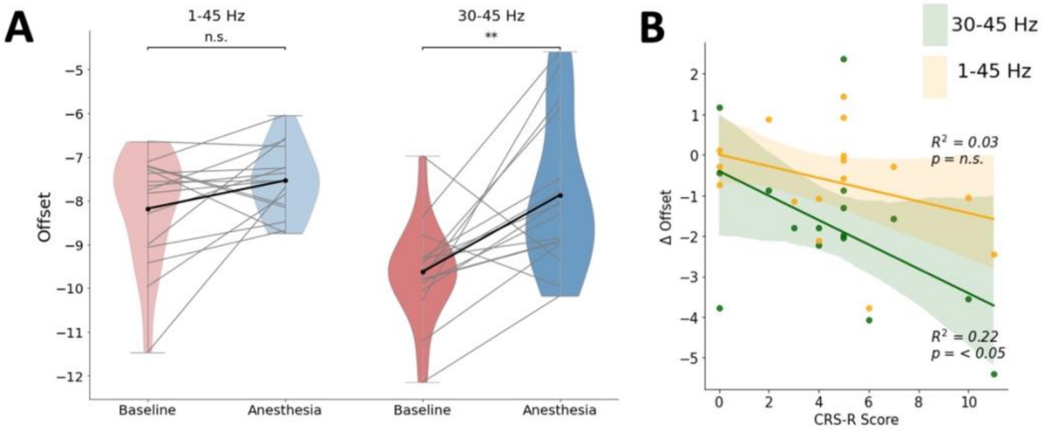


**Fig. S4.** Alterations of the spectral offset in DOC and general anesthesia. **(A)** Individuals offset significantly increased in response to propofol anesthesia in the 30-45 Hz range, but not the 1-45 Hz range. **(B)** The change in spectral offset in the 30-45 Hz range correlates with patients' CRS-R score.

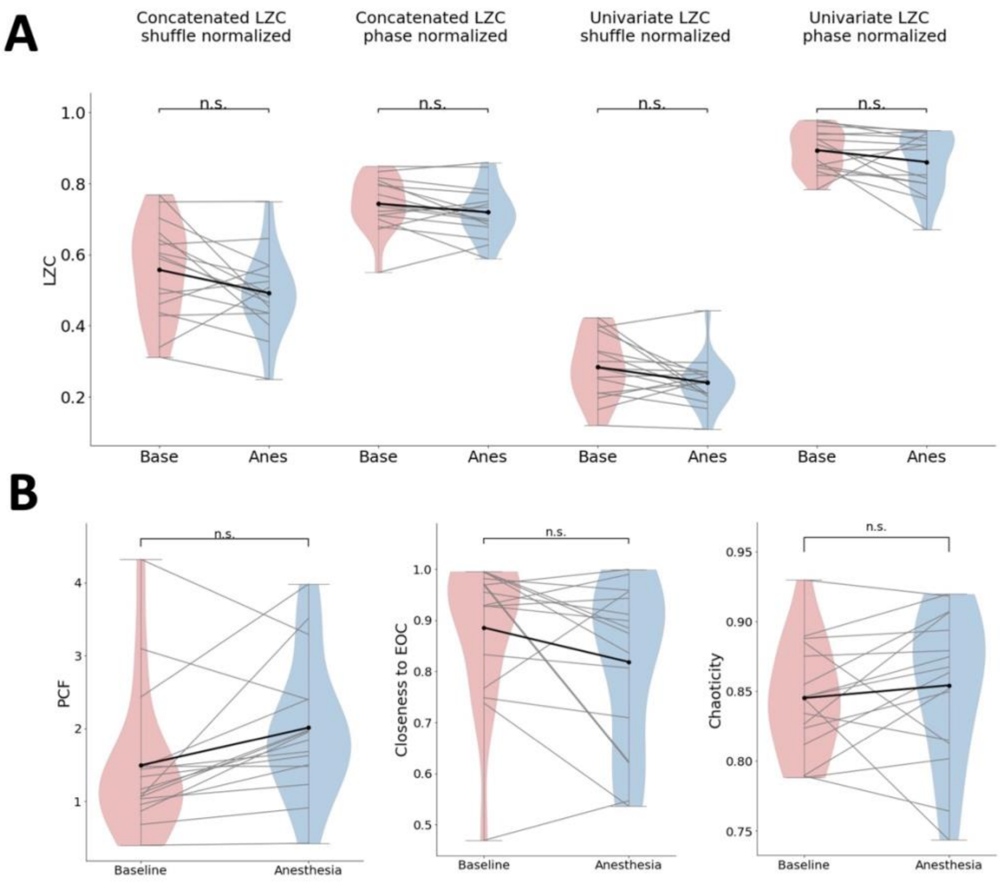


**Fig. S5.** Alterations of the network complexity and criticality in disorders of consciousness and general anesthesia. **(A)** Anesthetic-induced change of univariate and concatenated LZC, using two different normalization approaches. **(B)** Anesthetic-induced change of criticality, estimated by the pair correlation function (left) and closeness to the edge of chaos (middle). Anesthetic-induced change of chaoticity (right).

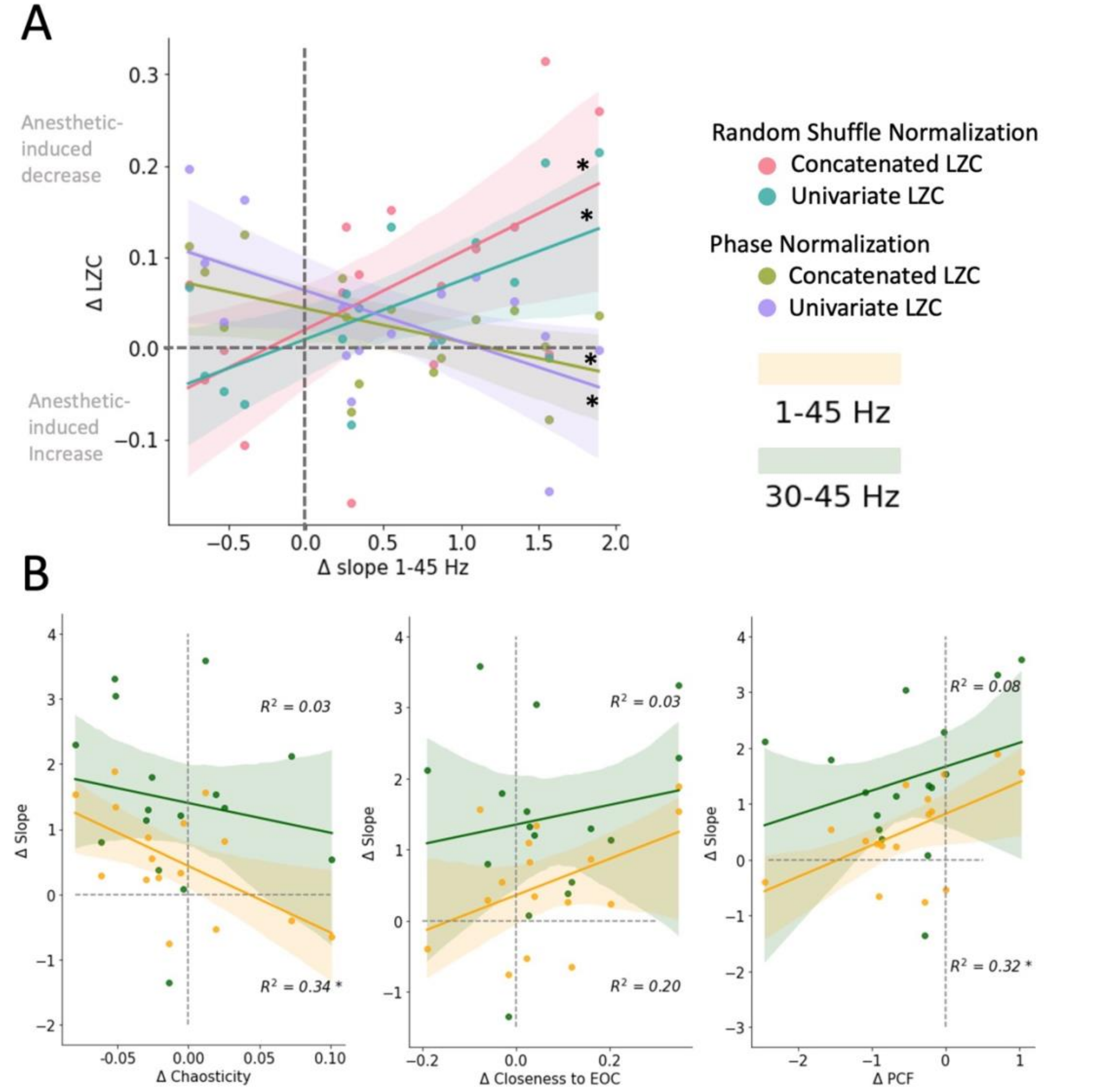


**Fig. S6.** Relation between the anesthetic-induced change of the spectral slope and the accompanying change of **(A)** signal complexity, using four different types of LZC, and ***(B)*** Chaoticity, Closeness to edge of chaos (EOC) and the pair correlation function (PCF). * indicates $p$<0.05, ** indicates $p$<0.01, *** indicates $p$<0.001.
